# Stability of mpox (monkeypox) virus in bodily fluids and wastewater

**DOI:** 10.1101/2023.05.09.540015

**Authors:** Claude Kwe Yinda, Dylan H. Morris, Robert J Fischer, Shane Gallogly, Zachary A. Weishampel, Julia R. Port, Trenton Bushmaker, Jonathan E. Schulz, Kyle Bibby, Neeltje van Doremalen, James O. Lloyd-Smith, Vincent J. Munster

**Affiliations:** Laboratory of Virology, Division of Intramural Research, National Institute of Allergy and Infectious Diseases, MT, USA; Department of Ecology and Evolutionary Biology, University of California, Los Angeles, United States; Department of Civil and Environmental Engineering and Earth Sciences, University of Notre Dame, Notre Dame, Indiana 46556, USA

**Keywords:** Mpox, human monkeypox virus, stability, surface, body fluid

## Abstract

**Importance:** Since May 2022, human monkeypox (mpox) infections have spread rapidly outside endemic countries. On July 23, 2023, WHO declared a Public Health Emergency of international Concern because of the unprecedented global spread of mpox. Whereas there is an incomplete understanding of transmission routes, the spread of monkeypox virus (MPXV) through sexual contact networks of men who have sex with men (MSM) highlights the potential of transmission during sexual activity, either via bodily fluids or via direct contact. This research assesses the potential for MPXV to remain infectious in the environment on surfaces and in fluids under different environmental conditions.

**Objective:** To investigate the stability of MPXV on various surfaces, at different environmental conditions and in different bodily fluids (human blood, semen, serum, saliva, urine, and feces), and to assess decontamination of MPXV-contaminated wastewater via chlorination.

**Design, Setting, and Exposures:** All experiments involving viable MPXV were performed at the Rocky Mountains Laboratory under high containment conditions using MPXV strain hMPXV/USA/MA001. Infectious MPXV was quantified via plaque assay. Environmental decay rates were estimated using a Bayesian statistical model, with a Poisson likelihood for plaque counts.

**Results:** MPXV showed surface dependent stability, with faster decay at higher temperatures. Decay rates varied considerably depending on the medium in which virus was suspended, both overall MPXV displayed considerable environmental stability in bodily fluids and in particular proteinaceous fluids, such as blood and semen, lead to greater persistence. Wastewater chlorination was an effective decontamination technique.

**Conclusions and Relevance:** MPXV retains its infectivity in the environment when it is deposited in blood, semen, and serum; environmental decay is more rapid in less proteinaceous fluids, particularly once residual liquid has evaporated. The observed persistence of MPXV implies environmental contamination in healthcare setting, could serve as vehicles of transmission and dissemination in these environments. Chlorination was effective in decontamination of wastewater. Key findings and public health implications; these results suggest that MPXV can have prolonged stability in the environment and are therefore directly relevant for hospital hygiene practices to prevent nosocomial transmission.

## Introduction

Human monkeypox (mpox) is a zoonotic infectious disease caused by monkeypox virus (MPXV). There are two known clades of MPXV: Clade I (formerly Congo basin clade) and Clade II (formerly West African clade)^1^. MPXV is endemic in Central and Western Africa, with periodic spillover into humans and limited onward transmission^2^. Historically, cases have been identified sporadically outside of endemic regions, mostly related to travel, nosocomial contact, or contact with imported rodents^3^. Beginning in May 2022, the largest known human outbreak of mpox occurred, caused by Clade IIb MPXV. On July 23 2022, the WHO declared the human monkeypox outbreak to be a public health emergency of international concern. Since then, over 87,000 laboratory confirmed cases have been reported, most outside of endemic regions. Human-to-human transmission of MPXV is likely to occur via direct contact transmission, or potentially by fomite transmission^3^. In the current outbreak, most cases involve men who have sex with men (MSM), with indications that sexual activity facilitates transmission. This can occur via skin-to-skin contact or through dissemination of bodily fluids. MPXV has been detected in a wide variety of samples including blood, saliva, urine, feces, semen, and skin, rectal and oropharyngeal swabs^4,5^. Environmental sampling detected low amounts of viable MPXV on household surfaces after even 15 days^6^. In addition, MPXV genetic material has been detected in wastewater streams^7^. All these have prompted concerns about nosocomial infection and risks to wastewater workers or possible reverse spillover into populations^8^. Here we evaluated the surface and body fluid stability of hMPXV/USA/MA001/2022, isolated in May 2022 from a human case in Massachusetts, USA and assess it wastewater decontamination potential using chlorination.

## Methods

All experiments were performed in triplicate for each surface and each environmental condition, starting with 10^5^ PFU/mL. Viable MPXV was quantified using plaque assays. Individual titers and virus half-lives were inferred in a Bayesian framework, modeling the plaque counts observed in titration wells as Poisson distributed (see the Methods section in the Appendix for details of the methods).

We measured decay of infectious MPXV in Dulbecco’s Modified Eagle Medium (DMEM) on multiple surfaces—cotton, stainless steel, and polypropylene—at multiple temperature/humidity conditions - 4°C, 45% RH, 21–23°C, 40% RH, and 28°C, 65% RH, and decay in human bodily fluids in bulk liquid solution and on polypropylene surface at 21–23°C.

To assess the effectiveness of chlorination as a decontamination technique, we measured virus decay in bulk deionized water and wastewater at multiple chlorine concentrations ranging from 0 to 10 parts per million (ppm).

## Results

In DMEM on surfaces, MPXV showed a biphasic decay pattern of initially slow decay followed by rapid decay. Since the transition typically occurred when all visible liquid had evaporated from the surface, we term these the wet phase and the dried phase. The virus was less stable at higher temperatures (Figure 1A). It was more stable on stainless-steel and polypropylene surfaces than on cotton, though note that recovery of viable virus from a porous, absorbent surface like cotton may be different from recovery from a non-porous, non-absorbent surface like stainless steel (Figure 1A). Posterior median estimated half-lives [2.5%, 97.5% quantile range] for the wet and dry phases are given in Supp. Table 1. A half-life on cotton for the dried phase could not be estimated for the 21–23°C and 28°C conditions, as no viable virus could be detected after the point of macroscopic drying (Figure 1A).

**Figure 1:**
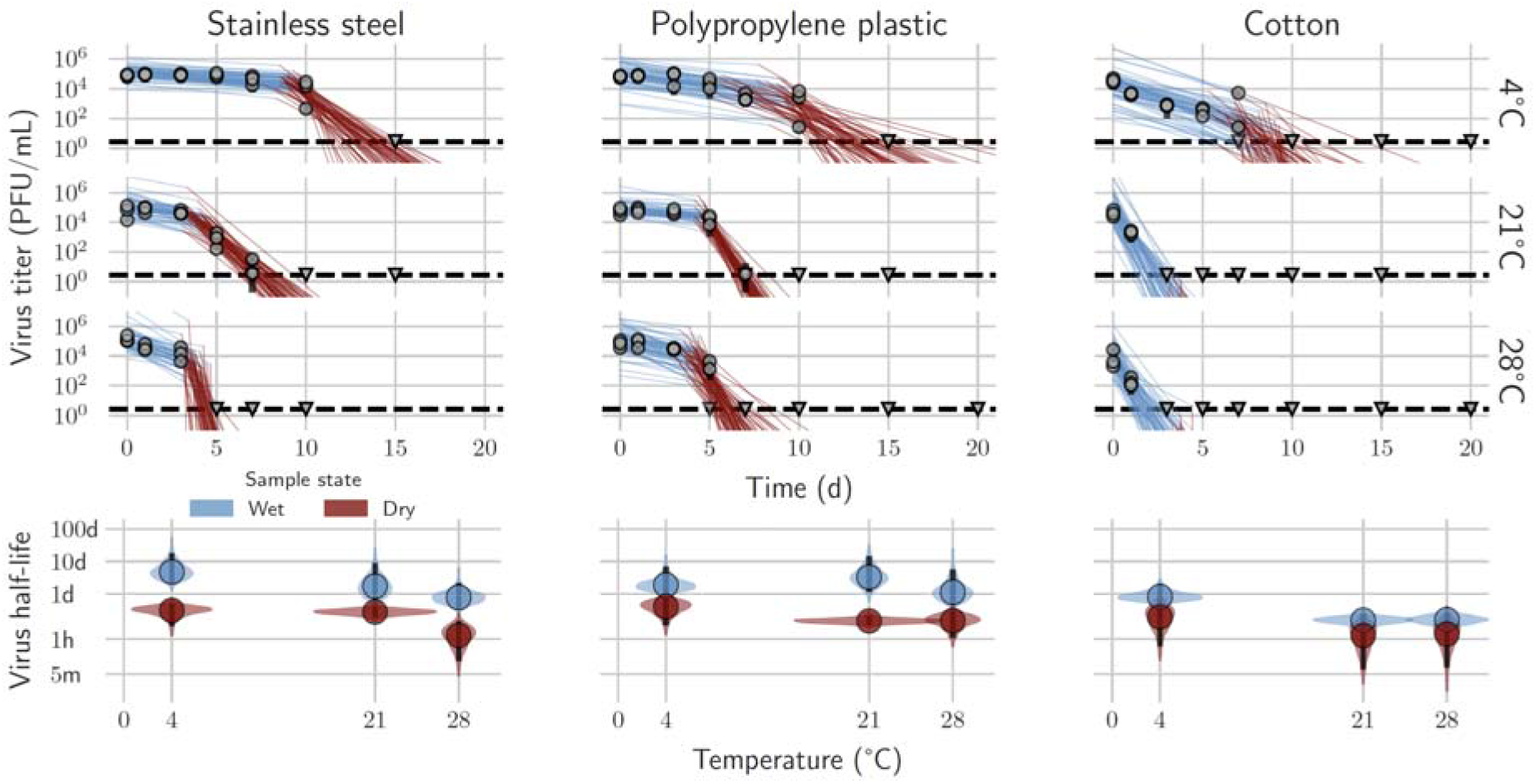
MPXV decay on cotton, polypropylene, and stainless steel at different environmental conditions. **A**. Regression lines showing predicted exponential decay of virus titer over time compared to measured (directly inferred) virus titers. Points show posterior median measured titers; black lines show a 95% credible interval. Colored lines are random draws from the joint posterior distribution of the exponential decay rate (negative of the slope) and intercept (initial virus titer); this visualizes the range of possible decay patterns for each experimental condition. Blue lines show the inferred “wet” phase, when visible residual moisture remains on the surface; red lines show the inferred “dried” phase, when evaporation has reached a state of quasi-equilibrium. The exact breakpoint is inferred from the data with a prior based on the last day of observable liquid. **B**: Inferred virus half-lives by surface and temperature condition. Violin plots show the shape of the posterior distribution. Dots show the posterior median half-life estimate and black lines show a 68% (thick) and 95% (thin) credible interval. Blue and red violin plots show inferred wet phase and dry phase half-lives, respectively.

Next, we investigated the stability of MPXV in human blood, semen, serum, saliva, urine, and feces (Figure 2A). All matrices were evaluated on surfaces as well as in bulk solution (“liquid”) form. Virus in blood, semen, and serum did not show an obvious difference in half-life between the wet phase and the dried phase, and both phases had half-lives similar to the half-life in bulk DMEM liquid (Figure 2B). In contrast, in saliva, urine, and feces, the virus showed a longer difference between a slower initial wet phase surface half-life, which for all secretions was again similar to the half-life in bulk liquid, and a substantially shorter half-life during the dried phase (Figure 2B).

**Figure 2:**
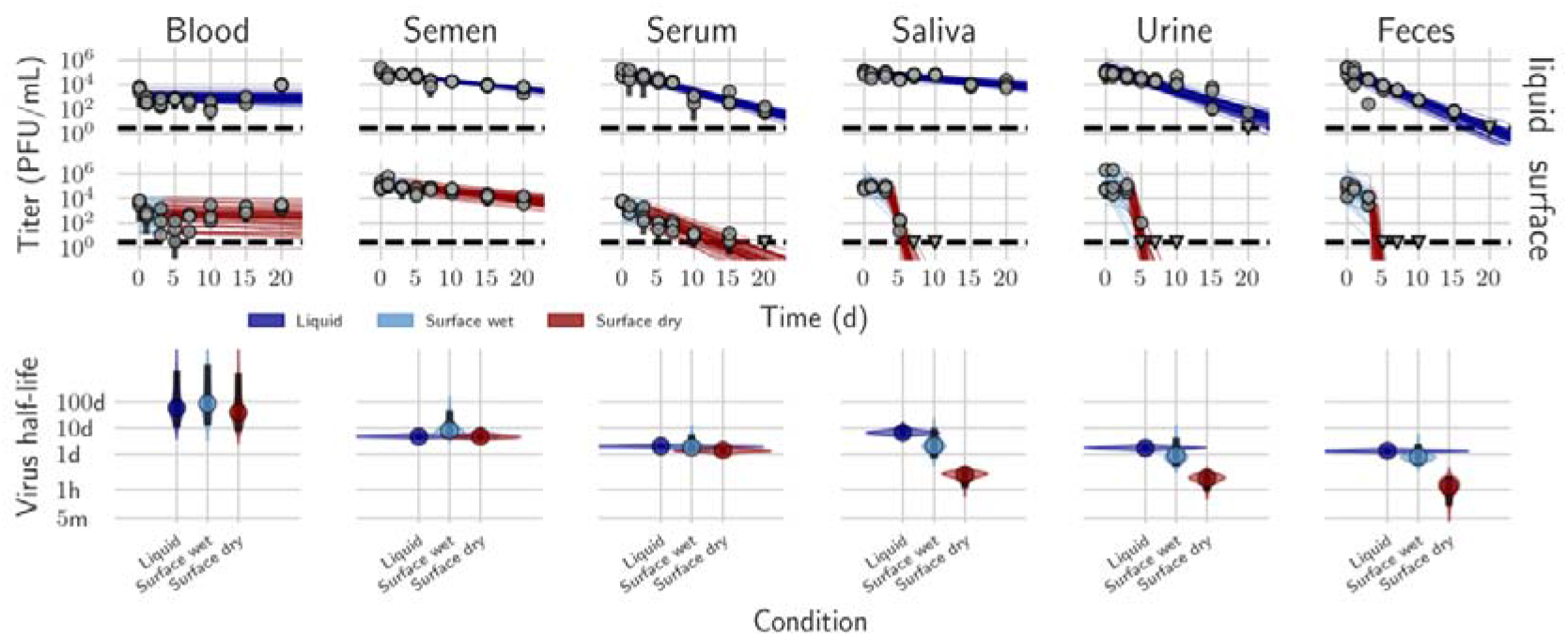
MPXV decay in human blood, semen, serum, saliva, urine, and feces on surface in liquid form. **A**. Regression lines showing predicted exponential decay of virus titer over time compared to measured (directly inferred) virus titers. Points show posterior median measured titers; black lines show a 95% credible interval. Colored lines are random draws from the joint posterior distribution of the exponential decay rate (negative of the slope) and intercept (initial virus titer); this visualizes the range of possible decay patterns for each experimental condition. Top row shows experiments in bulk solution (“liquid”); bottom row shows surface experiments. For surface experiments, light blue lines show the inferred “wet” phase, when visible residual moisture remains on the surface; red lines show the inferred “dried” phase, when evaporation has reached a state of quasi-equilibrium. The exact breakpoint is inferred from the data with a prior based on the last day of observable liquid. **B**. Inferred virus half-lives by condition and state. Violin plots show the shape of the posterior distribution. Dots show the posterior median half-life estimate and black lines show a 68% (thick) and 95% (thin) credible interval. Dark blue, light blue, and red violin plots show inferred bulk liquid, surface wet phase and surface dry phase half-lives, respectively.

For blood and semen, MPXV showed little or no detectable decay of infectious virus during the test period of 20 days (chosen arbitrarily), either in bulk liquid or on surfaces (Figure 2A, T_1/2_: blood liquid = 58.90 [10.00, 1638.42] days, blood surface dried phase = 38.75 [6.75, 1234.38] days, semen liquid = 4.63 [3.94, 5.70] days, semen surface dried phase = 4.57 [3.35, 7.09] days), though results for blood on surfaces were notably noisy. In serum, MPXV decayed over the test period, but with a long half-life of around a day or more (T_1/2_: liquid = 1.93 [1.71, 2.27] days, surface dried = 1.32 [0.98, 1.78] days) (Figure 2A,B).

MPXV had a long half-life in saliva both in bulk liquid and during the surface wet phase (Figure 2B, T_1/2_: liquid = 6.49 [4.72, 10.75] days, surface wet phase = 2.05 [0.66, 9.84] days), but it decayed rapidly during the dried phase (T_1/2_ surface dried phase = 0.16 [0.05, 0.25] days). The virus was less stable in urine and feces, but again with accelerated decay during the surface dried phase (Figure 2B). For urine, half-lives were 1.69 [1.35, 2.17] days in bulk liquid, 0.86 [0.32, 4.10] days during the surface wet phase, and 0.11 [0.03, 0.21] days during the surface dried phase. For feces, half-lives were 1.28 [1.07, 1.53] days in bulk liquid, 0.76 [0.35, 2.51] days during the surface wet phase, and 0.06 [0.01, 0.14] days during the surface dried phase.

Overall MPXV was consistently at least as stable in bulk liquid environments compared to surface conditions, particularly wet surface conditions. Major wet versus dry differences were apparent for saliva, urine, and feces, but not blood, semen, and serum. Based on these differences in decay patterns and on prior work on other viruses^9^, we hypothesized that a highly proteinaceous environment provides protection against decay of the virus, perhaps particularly during and after the evaporation of residual water following deposition. To investigate this, we assessed the stability of the virus in a bulk solution consisting of increasing amounts of serum (0%, 40%, 80%, 100%) mixed with DMEM. We saw monotonically increasing virus stability (measured as an increased half-life) as a function of the percentage of serum (Supplementary Figure 1).

Lastly, to assess the need for and effectiveness of wastewater decontamination against MPXV, we determined the stability and sodium hypochlorite inactivation of MPXV in wastewater and deionized (DI) water (Figure 3). In untreated DI water, MPXV did not show decay during the sampling period: (T_1/2_ = 60.79 [22.67, 1078.62] days, Figure 3). There was meaningful decay in wastewater, but with a long half-life of multiple days (T_1/2_ = 5.74 [4.58, 8.05] days, Figure 3).

**Figure 3:**
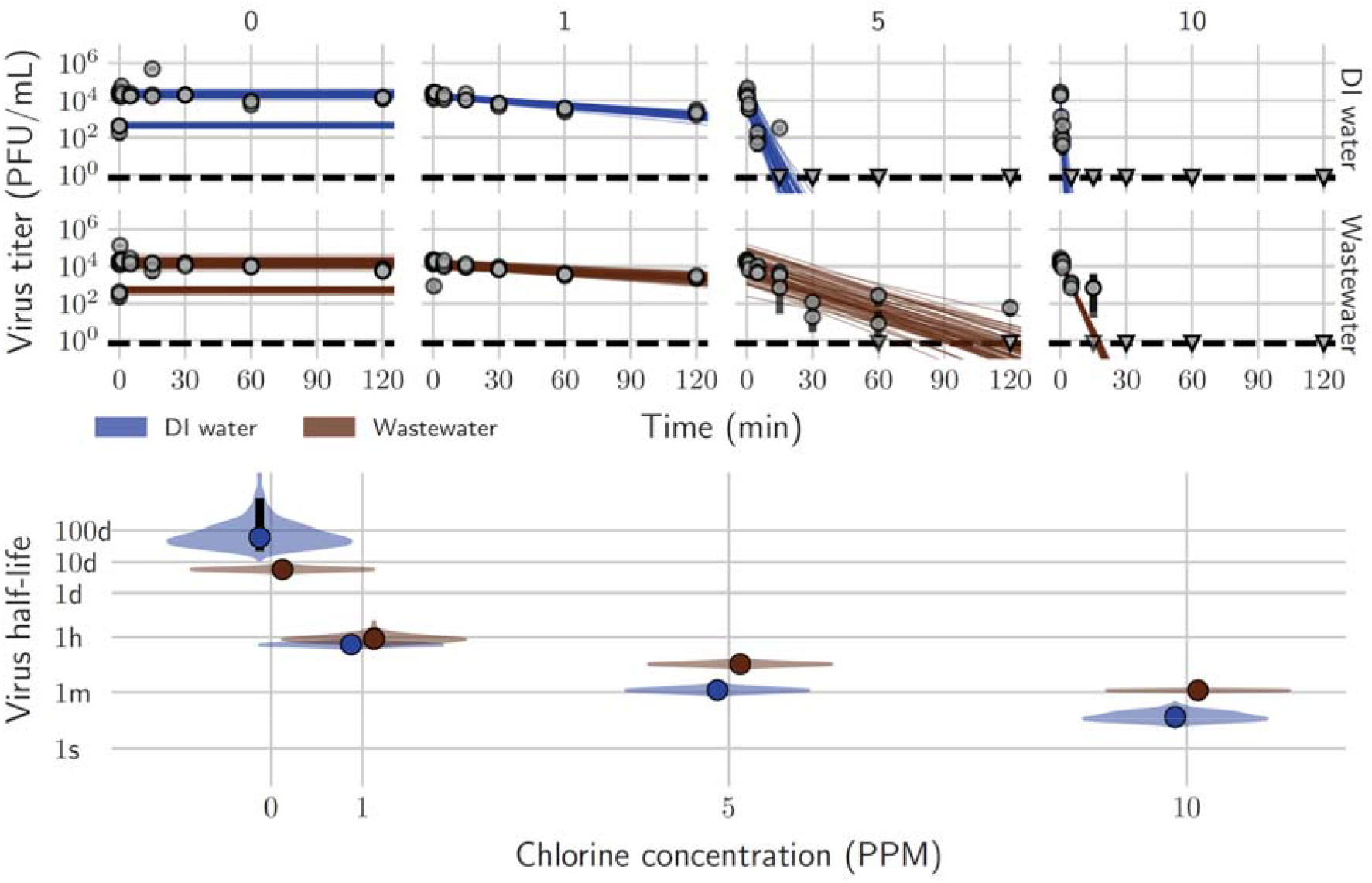
MPXV exponential decay and decontamination in wastewater and deionized water. **A**. Regression lines showing predicted decay of virus titer over time compared to measured (directly inferred) virus titers. Points show posterior median measured titers; black lines show a 95% credible interval. Colored lines are random draws from the joint posterior distribution of the exponential decay rate (negative of the slope) and intercept (initial virus titer); this visualizes the range of possible decay patterns for each experimental condition. **B**. Inferred virus half-lives as a function of free chlorine concentration in ppm. Violin plots show the shape of the posterior distribution. Dots show the posterior median half-life estimate and black lines show a 68% (thick) and 95% (thin) credible interval.

For sodium hypochlorite inactivation, rapid disinfection of MPXV was observed in DI water, with half-lives of 1.19 [0.85, 1.71] minutes at 5 ppm free chlorine and 0.17 [0.10, 0.34] minutes at 10 ppm. In wastewater, higher concentrations were required for rapid inactivation of highly contaminated samples: the half-life of viable virus was 8.13 [6.45, 10.50] minutes at 5 ppm and 1.17 [1.05, 1.28] minutes at 10 ppm. Differences in required chlorine concentrations could be due to high free chlorine consumption by the wastewater^10^. These results suggest that MPXV is quite stable in water, including wastewater, but that wastewater disinfection could quickly and significantly reduce the levels of viable virus.

## Discussion

Few studies have investigated the stability of viruses in the genus Orthopoxvirus (family *Poxiviridae*). Prolonged smallpox stability has been reported in scabs, vesicle and pustule fluids, lymph, and purulence of patients; variola in raw cotton and vaccinia in storm water^11-13^. The only experimental study on MPXV (Clade I) was conducted in aerosol and indicated that MPXV could remain viable in aerosol form for a prolonged period^14^.

Here, we found that MPXV indeed shows strong environmental persistence on surfaces and in bulk liquid. MPXV showed clear biphasic decay on surfaces for some media (DMEM and authentic human saliva, urine, and feces), but not others (blood, semen, and serum). The observed biphasic decay is indicative of different stability kinetics when the virus is initially deposited in a liquid solution versus after macroscopic water has evaporated.

In the different evaluated fluids and wastewater, the stability was fluid-type dependent. The persistence of MPXV in clinical specimens^15^ or tissues also suggests a fluid-type dependent decay. More proteinaceous solutions such as blood, serum, and semen favored virus stability; we confirmed experimentally that the protective effect of serum was concentration-dependent. This is consistent with other recent observations that protein in solution can thwart virus environmental inactivation, and that virus environmental inactivation rate can depend strongly on physicochemical properties of the medium^9^.

Environmental risk assessment has typically focused on properties of the ambient environment (temperature, humidity, surface type for fomite transmission, ventilation rate for airborne transmission). Taken together with prior work on other viruses, our results suggest that route and type of contamination may also be important to consider, as viruses may be much more or less viable depending on the human secretions in which they are shed. This may partly account for the variability observed in the environmental contamination of MPXV on different types of household surfaces^6^. In addition, this explains the discrepancy between the longevity of MPXV on cotton as evaluated in this study compared to the reported results from epidemiological investigations where exudate (vesicular or pustular fluids) will provide a more protective environment^12^. Nevertheless, the persistence of MPXV in the environment suggest that necessary precautions are required to avoid environmental transmission and particularly nosocomial transmission in hospital settings.

Sexual transmission of MPXV has been confirmed given the virus’s spread through sexual contact networks of MSM. The long half-life of viable MPXV in blood and semen increase the transmission risk via fluid exchange, though it is important to note that transmission via skin-to-skin contact during sexual activity is also plausible. The potential for long-term viability in semen has further implications for the viral load in semen within an infected individual, and for the duration of infectiousness after viral replication stops. The half-lives estimated in our work are consistent with infectious virus remaining present in semen for weeks after virus replication ends.

Our finding that MPXV can remain infectious for weeks in untreated wastewater raises potential for exposure of sanitation workers, peridomestic animals, and wildlife^8^. Given the suspected role of rodents as reservoirs of MPXV, this raises hypothetical concerns about establishment of zoonotic reservoirs in previously non-endemic countries. However, we emphasize that dilution will mitigate these risks. Our findings suggest that testing for infectious MPXV could be a valuable complement to PCR-based wastewater surveillance when significant quantities of viral DNA are detected.

## Conclusion

The results suggest that MPXV stability is dependent on the surface, environmental conditions, and the matrix of the virus. Overall, MPXV showed long half-lives in a variety of human fluids, both in bulk liquid solution and when deposited on common surfaces; we also found half-lives approaching a week in untreated wastewater. This suggests that virus stability is sufficient to support environmental or fomite transmission of MPXV, so exposure and dose-response will be the limiting factors for these transmission routes.

## Acknowledgments

We would like to thank Zachary Wiener, Todd Smith, Nicolle Baird, Christina Hutson, Fahim Atif, and Inger Damon of the CDC for rapidly sharing the MPXV strain used in this study. We would also like to thank Bernie Moss, Patricia Earl, Elaine Haddock, the RML institutional biosafety committee, and the biosafety office for helpful suggestions and support.

