## appendix for "Stability of mpox (monkeypox) virus in bodily fluids and wastewater"

**Methods**

We performed all experiments involving viable MPXV at Rocky Mountain Laboratories (RML) under BSL4 condition using MPXV strain hMPXV/USA/MA001/2022 (4.8 × 10^6^ PFU/mL, hereafter called MA001) (BEI Resources). We propagated the virus in VeroE6 cells in Dulbecco’s modified Eagle’s medium (Sigma-Aldrich, St, Louis, MO) supplemented with 10% fetal bovine serum, 1 mM L-glutamine, 50 U/mL penicillin and 50 μg/mL streptomycin (10% DMEM). All experiments were completed in triplicate at room temperature (21 – 23 °C) unless otherwise indicated. MPXV was quantified using a plaque assay. The limit of detection for all replicates was 2 PFU/mL. All experimental measurements are reported as median across three replicates. Human body fluids were commercially acquired from Lee BioSolutions Inc. (https://www.leebio.com). Wastewater samples were collected from a municipal wastewater treatment plant in northern Indiana, United States, then shipped frozen overnight to RML where they were stored at -80 °C until used as previously described^1^.

*MPX-V stability on surfaces at different environmental conditions*

The surface stability of MPXV MA001 was evaluated on 15 mm polypropylene and AISI 316L alloy stainless steel discs at 4°C/40%RH, 21-23°C/40% RH, and 28°C/65%RH. 50 μL of MPXV stock containing 10^5^ PFU was deposited (7-10 drops) on the surface of a disc. At predefined time-points (0, 1, 3, 5, 7, 10, 15, 20 days after deposition) viable virus was recovered by rinsing with 1 mL of Dulbecco’s modified Eagle’s medium (Sigma-Aldrich, St, Louis, MO) supplemented with 2% fetal bovine serum, 1 mM L-glutamine, 50 U/ml penicillin and 50 μg/ml streptomycin (2% DMEM) and frozen at -80°C until titrated.

*MPX-V stability in human body fluid*

Stability was measured on the surface by pipetting 50 μL of each fluid containing 10^5^ PFU of MPXV on plastic or left in a bulk liquid containing 2.0 × 10^6^ PFU/mL (10^5^ PFU/50 μL, stored in a screw-top vial) at 21°C/45%RH.

To determine the stability of MPXV in body fluids, human saliva, semen, fecal matter, and urine were spiked with MPXV MA001. To determine the stability of the virus in body fluids deposited on a surface and allowed to dry naturally, 50 μL of each fluid containing 10^5^ PFU of virus was aliquoted onto a polypropylene disc and left at room temperature (21 – 23°C) with 40% RH. Sample recovery was performed at predefined time-points (0, 1, 3, 5, 7, 10, 15, 20 days after deposition) by rinsing with 1 mL of 2% DMEM and frozen at -80 °C until titrated. To determine the virus stability in a bulk liquid, we initially prepared fluid containing 2.0 × 10^6^ PFU/mL (10^5^ PFU/50 μL). The bulk fluid samples were stored in a screw-top vial at room temperature between sampling times. At 0, 1, 3, 5, 7, 10, 15, 20 days after deposition 50 μL of each fluid-virus mix was pipetted into 1 mL of 2% DMEM and frozen at -80 °C until titrated.

*MPXV stability in wastewater and DI water*

50 µL of stock virus was diluted in 5 mL of wastewater or DI water (1:100 dilution) and sample recovery was performed at predefined time-points (0, 1, 3, 5, 7, 10, 15, 20 days after deposition) by rinsing with 1 mL of 2% DMEM and frozen at -80 °C.

To assess the stability of MPXV in wastewater and in DI water 50 µL of stock virus was diluted in 5 mL of wastewater or DI water (1:100 dilution) in triplicate. At time 0 and at 1-, 3-, 5-, 10-, 15-, and 20-days 100 µL of virus spiked sample was put into 900 µL of Dulbecco’s modified Eagle’s medium (Sigma-Aldrich, St, Louis, MO) supplemented with 2% fetal bovine serum, 1 mM L-glutamine, 50 U/ml penicillin and 50 μg/ml streptomycin (2% DMEM) and frozen at -80°C until titrated. The physiochemical parameters of the wastewater have been previously reported ^1^

*Wastewater disinfection*

To test the efficacy of free chlorine for wastewater disinfection of MPXV, stock virus was diluted 100X in wastewater or DI water and 1.098 mL was added to the top row of a deep well 96-well plate. The ‘time zero’ sample was taken prior to the addition of chlorine to obtain the initial virus concentration in the sample. Sodium hypochlorite (Acros Organics, Fair Lawn, NJ) was added to obtain an initial dose of 0, 1, 5, or 10 parts per million (ppm) to each of three wells. At 0, 1, 5-, 10-, 30-, and 60-minute time points, 100 µL samples were added to 100 µL of 0, 1, 5, or 10 ppm sodium thiosulfate solution to quench remaining free chlorine. The resulting solution was triturated and transferred in total to 800 µL of DMEM supplemented with 2% fetal bovine serum, 1 mM L-glutamine, 50 U/ml penicillin and 50 μg/ml streptomycin. These samples were frozen at -80°C until titrated.

*Virus quantification via endpoint titration plaque assay*

Frozen samples were thawed and a 10X serial dilution was performed. 250 µL of each dilution was added to a well of confluent Vero E6 cells in a 12-well plate and incubated for 2 hours. After 2 hours an additional 1 mL of 2% DMEM was added to each well. The plates were incubated at 37°C with 5% CO_2_ for 4 days. On day 4 the medium was removed from the wells and replaced with 10% formaldehyde for 10 minutes. At the end of 10 minutes the formalin was removed and replaced with a 1% solution of crystal violet. The crystal violet remained on the cells for 10 minutes at which point it was removed and the plates rinsed with water. After drying the plates were assessed for plaques.

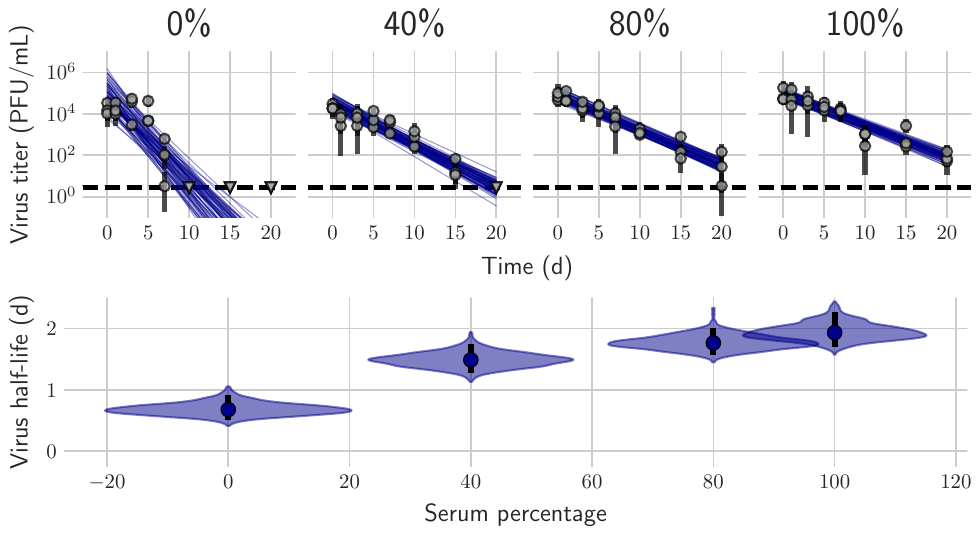
**Supplementary figure 1:** MPXV decay in different human serum dilutions in DMEM. **A**. Regression lines showing predicted exponential decay of virus titer over time compared to measured (directly inferred) virus titers. Points show posterior median measured titers; black lines show a 95% credible interval. Colored lines are random draws from the joint posterior distribution of the exponential decay rate (negative of the slope) and intercept (initial virus titer); this visualizes the range of possible decay patterns for each experimental condition. **B**. Inferred virus half-lives by condition and state. Violin plots show the shape of the posterior distribution. Dots show the posterior median half-life estimate and black lines show a 68% (thick) and 95% (thin) credible interval. Violins show half-lives.

*Reference*

1.	Bivins A, Greaves J, Fischer R, et al. Persistence of SARS-CoV-2 in Water and Wastewater. *Environmental Science &amp; Technology Letters*. 2020;7(12):937-942. doi:10.1021/acs.estlett.0c00730
